# *APOE ε4* allele advances the age-dependent decline of amyloid β clearance in the human cortex

**DOI:** 10.1101/2021.04.07.438832

**Authors:** Atsushi Saito, Yusuke Kageyama, Olga Pletnikova, Gay L. Rudow, Yang An, Yumi Irie, Akiko Kita, Kunio Miki, Ling Li, Pamela Southall, Kazuhiro Irie, Juan C. Troncoso

## Abstract

**Introduction:** Our previous study indicated that the pericapillary clearance of amyloid β (Aβ) declines with age in *APOE 3/3* subjects. Here, we examine whether the *APOE* ε*4* allele has an impact on this age-related decline.

**Methods:** We examined 69 autopsy brains of *APOE* ε*3/*ε*4* or *APOE* ε*3/*ε*3* individuals (30-65 years) for the immunohistochemical localization of intracellular, extracellular, and pericapillary Aβ in the cerebral cortex.

**Results:** In *APOE* ε*3/*ε*4* individuals, the percentage of Aβ positive pericapillary spaces began to decrease (p=0.030), and the number of extracellular Aβ particles increased in the early 30s (p=0.0008). Those average values were significantly lower (p<0.0001) and higher (p<0.0001), respectively, compared to *APOE* ε*3/*ε*3* individuals.

**Discussion:** Our observations indicate that *APOE* ε*4* allele advances by one decade at the onset of age-related decline in Aβ glymphatic clearance. This finding supports early clinical intervention and stratification by APOE genotype to prevent Aβ deposition and AD progression.

## 1 INTRODUCTION

Late onset or sporadic Alzheimer’s disease (AD) is the most common cause of dementia in the elderly population and its major risk factors are aging^1^ and the *APOE* ε*4* allele^2^. The pathological features of AD include amyloid β (Aβ) plaques, tau-based neurofibrillary tangles, inflammation^3^, and synaptic degeneration. Aβ is a peptide important for synaptic function and homeostasis^4^ and its aberrant accumulation leads to formation of Aβ plaques and subsequent neurodegeneration^5^. Aβ is constantly produced and cleared from the brain via multiple pathways including the glymphatic system^5^ and a decline of its clearance in aging has been implicated in the formation of Aβ plaques^6^. Previous pathological observations and imaging studies indicate that the onset of Aβ deposition in the cerebral cortex begins as early as the fifth decade of life^7^ and precedes the cognitive impairment of AD by decades^8^. However, the dynamics and clearance of Aβ before its deposition in plaques has not been fully assessed. Examination of Aβ pathophysiology before the onset of Aβ plaques is critical for the development of preventive interventions.

Our previous study of Aβ clearance focused on the pericapillary space, a compartment of the glymphatic system, in the brain of younger individuals with *APOE* ε*3/*ε*3* genotypes and showed an age-dependent decline in clearance with onset at approximately 45 years of age^9^, an observation consistent with aging as a major risk for AD^1^. The *APOE* ε*4* allele represents the other major risk factor for AD and there is strong evidence that this allele enhances Aβ deposition^10^ and a recent study associates *APOE* ε*4* with impaired protein clearance^11^. These results suggest that the *APOE* ε*4* allele would render the aging brain more susceptible to Aβ accumulation due to decline in clearance.

Here, we test the hypothesis that the *APOE*ε*4* allele advances the age-dependent decline of Aβ clearance via the glymphatic system. To this end, we first conducted additional biophysical and immunological characterization of the 11A1 antibody. Second, we used immunohistochemistry (IHC) to examine the dynamics of Aβ in the cerebral cortex and its clearance through the pericapillary spaces in the postmortem brains of *APOE* ε4 carriers 34 to 65 years of age, before the appearance of Aβ plaques. Based on these observations, we established the onset of decline of Aβ clearance and compared it with our dataset from an *APOE* ε*3/*ε*3* cohort^9^. Third, we assessed the size and number of extracellular Aβ aggregates (≥3µm) across the age spectrum in both cohorts.

## 2 METHODS

Details of all the experiments are described in e-Methods (supplementary information).

### 2.1 Autopsies, Subjects and Neuropathological examination

Autopsies of 34 subjects, ages 34-65 years, 65% males and 25% females (**Table 1 and Table A. 1**) with *APOE* ε*3/*ε*4*, and age, gender, and race matched those of 35 subjects with *APOE* ε*3/*ε*3* ^9^ (**Table 1**) were performed at the Office of the Chief Medical Examiner (OCME) of the State of Maryland in Baltimore, and brains were accessioned as previously described^9^. We followed protocols authorized by the Institutional Review Board (IRB) of the State of Maryland Department of Health and Human Services and Johns Hopkins Medicine.

**Table 1:** Sample characteristics

| | *APOE* $\epsilon 3/\epsilon 4$ | *APOE* $\epsilon 3/\epsilon 3$ | p value |
| --- | --- | --- | --- |
| N | 34 | 35 | - |
| Age<br>Mean (SD) | 48.3 (7.9) | 49.2 (10.7) | 0.67 |
| Sex<br>(Male%) | 64.7 | 65.7 | >0.99 |
| Race<br>(Caucasian%) | 76.5 | 77.1 | >0.99 |

Age was examined by t-test, Sex and race were examined by Fisher's exact test

### 2.2 Crystallographic and immunological characterization of the 11A1 antibody

To identify the recognition sites of 11A1 on the Aβ peptide, we performed X-ray crystal structure analysis of 11A1 Fab-hapten peptide (E22P-Aβ10-34). Validations of the immunoreactivity of 11A1 antibody to Aβ in human cortex was conducted by immunohistochemistry.

### 2.3 Evaluation of Aβ localization in inferior parietal cortex

We analyzed the distribution of 11A1 immunoreactivity (IR) in formalin-fixed paraffin embedded tissues of the inferior parietal cortex of *APOE* ε*3/*ε*4* individuals with the same strategy of the previous study of *APOE* ε*3/*ε*3* subjects^9^. We selected 34 cases (**Table 1 and Table A. 1**) with brain samples screened for “optimal tissue preservation” by assessing the integrity of cerebellar Purkinje cells on tubulin immunostains^9^. Each section was co-stained with 11A1 antibody and one of the following molecular markers: Microtubule Associated Protein 2 (MAP2) for neurons, Aldehyde Dehydrogenase 1 Family Member L1 (ALDH1L1) for protoplasmic astrocytes, Collagen IV perivascular spaces, or CD63 for extracellular vesicles.

A single operator counted the 11A1 IR and cell-type markers IR blinded to age, sex, race, and genotype.

### 2.4 Antibodies

The information of primary and secondary antibodies used in this study is listed in **Tables A. 2 and A. 3.**

### 2.5 Statistical analyses

We used Pearson correlations to quantify the linear relationship between age and distribution of IR. To characterize further the relationship of parameters with age, we compared three regression models with increasing complexities. The first model is the intercept only model, which means there is no age relationship. The second model is the intercept and age model, which assumes linear age relationship. The third model is the piecewise regression model with knot at age 45 years based on our previous study^9^, which assumes that age relationships are different before and after age 45 years. We compared the models using Akaike information criterion (AIC) (**Table A. 4**). Statistical analyses were performed with R version 3.4.4.

## 3 RESULTS

### 3.1 11A1 Antibody characterization

After extensive screening, we obtained crystals diffracting to 1.75 Å resolution (**Table A. 5**). As shown in **Figure 1A-C**, 11A1 Fab bound to Tyr10 to His14 region of E22P-Aβ10-34, indicating that 11A1 is an antibody to the N-terminal region, and does not recognize the positions 22 and 23 (toxic turn) directly as previously expected^12^. Notably, there are no previously reported Aβ antibodies that recognize the Try10-His14 epitope. For example, the epitopes of 82E1, 4G8, and Crenezumab are Asp1-Glu3, Leu17-Val24, and His13-Val24, respectively. There are various truncated Aβ species in the N-terminal region, which accumulate as intracellular aggregates (oligomers) and in senile plaques. Systematic proline replacement by Irie and colleagues^13^ suggested that the N-terminal 14 residues did not form an ordered structure, and that the residues 15-21 were involved in the intermolecular β-sheet in the aggregates (oligomers). Because 11A1 recognizes the disordered Tyr10-His14 region, which is adjacent to the core region of Aβ aggregates with intermolecular β-sheet structure (Gln15-Ala20), this antibody could detect most of the truncated species of Aβ as well as native Aβ with high sensitivity.

**Figure 1.**
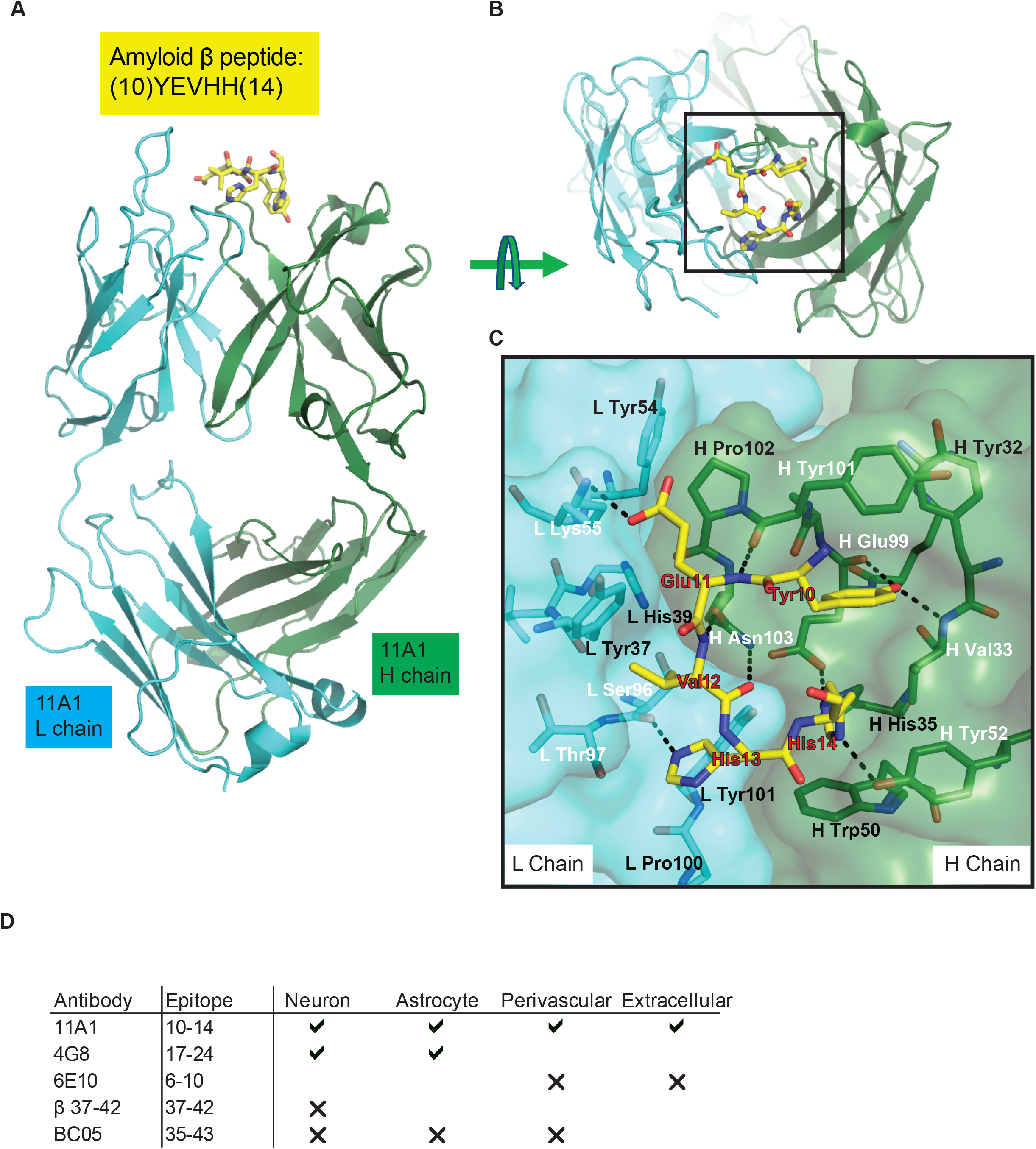
Validation of 11A1 antibody. Crystal structure of 11A1 Fab in complex with Aβ10-34 **(A)** front view, **(B)** top view, **(C)** interactions among amino acid residues of 11A1 and Aβ10-34. The amino acid residues at positions 10-14 of Aβ10-34 are recognized by 11A1. **(D)** Result of Immunofluorescent staining with commercial Aβ antibodies. Details of the antibodies are shown in **Table A. 2** and **A. 3**. Check mark: Signal found, sufficient signal noise ratio (SNR) for analysis, Blank: Signal found, not sufficient SNR for analysis, X mark: Signal not found.

### 3.2 Comparison of 11A1 antibody to other Aβ antibodies on immunostaining

Next, we compared the immunoreactivity (IR) of 11A1 to that of other Aβ antibodies that target different epitope sequences than 11A1. We tested commercially available antibodies 4G8, 6E10, BC05 and C-term. Among these, 4G8 showed the closest IR pattern to 11A1. All of the antibodies recognize Aβ plaques, but show some differences in the recognition of Aβ in specific cells (**Figure A. 1**). As shown in **Figure 1D**, 11A1 appears as the best antibody for this study based on its ability to recognize Aβ in multiple cellular and extracellular compartments, as well as perivascular spaces. We found that 11A1 is able to detect a broader type of Aβ distribution than the other commonly used Aβ antibodies such as 6E10 and 4G8. The results in the crystallographic study and the comparison of 11A1 IR with other Aβ antibodies strengthen the relevance of 11A1 IR to studies of Aβ in general.

### 3.3 Quantitation of 11A1 immunoreactivity in neurons, astrocytes, pericapillary spaces, and the extracellular compartment of the inferior parietal cortex

First, we determined the percentage of neurons displaying 11A1 colocalization signal in their perikaryon among inferior parietal neurons characterized by MAP2 labeling (**Figure 2A**). Similar to observations on *APOE* ε*3/*ε*3* individuals^9^, 11A1 IR was present in a high percentage (85%) of neurons across the age spectrum. Regression analyses showed that the intercept model fit the data best (using Akaike information criterion [AIC], **Table A. 4**), meaning that the percentage of labeled neurons did not show a significant relationship with age (p=0.26, **Figure 2B**).

**Figure 2.**
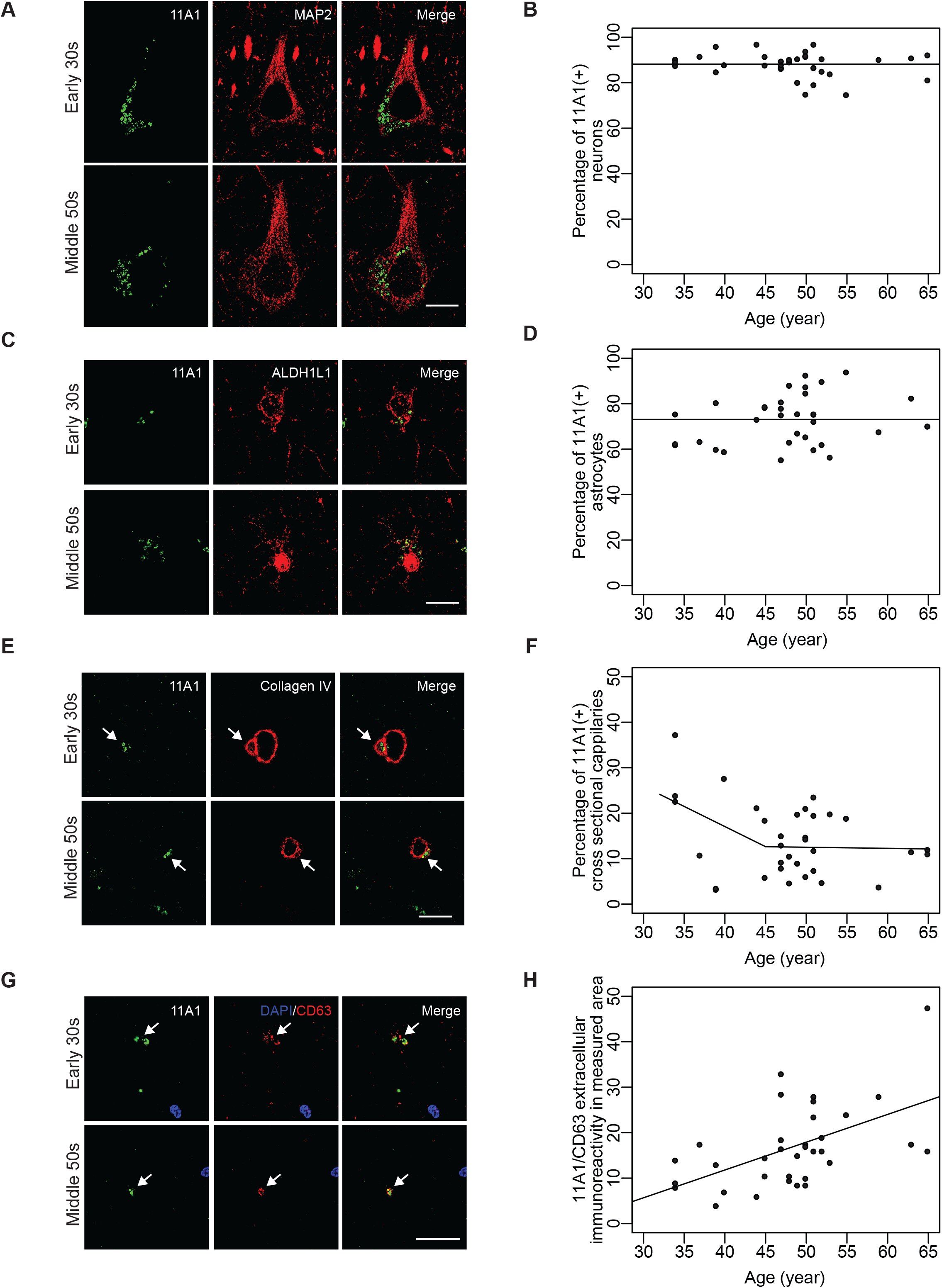
11A1 immunoreactivity in *APOE* ε*3/*ε*4* individuals. (A, C, E, G) Representative images of 11A1 co-stained with the molecular markers; **(A)** MAP2, **(C)** ALDH1L1, **(E)** Collagen IV, **(G)** CD63, respectively. The rows of Figures correspond to two age groups: early 30s and middle 50s. Left panels show the 11A1 signal, middle panels indicate the molecular markers signal and right panels show merged images. *Scale bars*, 5µm. **(B)** Percentage of cortical neurons (MAP2 positive cells) labeled with 11A1 in measured area (0.910 mm^2^) across the age-spectrum (34-65 years). The relationship with age is not significant (p=0.20). **(D)** Percentage of cortical astrocytes (ALDH1L1 positive cells) labeled with 11A1 in measured area across the age-spectrum (34-65 years). The relationship with age is not significant (p=0.13). **(F)** Percentage of perivascular spaces labeled with 11A1 in measured area across age-spectrum (34-65 years). The relationship with age is significant by age of 45 (p=0.030), is not significant after age of 45 (p=0.92). **(H)** Number of immunoreactivity of extracellular 11A1 particles (≤ 1µm) in measured area (0.407mm^2^) across age-spectrum (34-65 years). The relationship with age is significant (p=0.0084).

Second, we examined 11A1 IR in protoplasmic astrocytes co-stained with ALDH1L1 (**Figure 2C**). 11A1 IR was present in 60% to 90% of astrocytes across the age range. We found the intercept model fit the data best (**Table A. 4**), indicating that the percentage of 11A1 IR positive astrocytes did not show a significant relationship with age (p=0.19, **Figure 2D**).

Third, we examined pericapillary spaces identified with Collagen IV. Similar to the observation in the *APOE* ε*3/*ε*3* individuals^9^, 11A1 immunoreactive particles were present in the pericapillary spaces across the age span (**Figure 2E**). Regression analyses showed that the piecewise age model with knot at age 45 fit the data best (using Akaike information criterion [AIC], **Table A. 4**). The percentage of pericapillary spaces labeled with 11A1 IR tended to decrease linearly before age 45 years (p=0.030, **Figure 2F**), but remained stable thereafter (p=0.92, **Figure 2F**). Lastly, we counted extracellular particles, displaying 11A1 and CD63 colocalization, measuring <1µm in diameter, and located at a distance ≥5µm from the edge of DAPI signal (nuclear marker) (**Figure 2G**) as described in eMethods. The number of extracellular 11A1/CD63 particles per area increased significantly throughout the age range (p=0.0008) (**Figure 2H**) and the linear regression model fit best for its relationship with age (using Akaike information criterion [AIC]) (**Table A. 4**).

### 3.4 Comparison between *APOE* ε*3/*ε*3* and *APOE* ε*3/*ε*4* individuals

Our previous studies indicate that the presence of the *APOE* ε*4* allele accelerates Aβ plaque formation^7,14^. Thus, we examined whether the dynamic of 11A1 IR in the various brain compartments is different in *APOE* ε*3/*ε*4* and *APOE* ε*3/*ε*3* individuals^9^ before Aβ plaques form. Comparing the two genotypes, we observed no difference in the percentage (p=0.057, **Figure 3A**) or in the relationship with age (p=0.34, **Figure 3B**) of 11A1-labeled neurons, or those of astrocytes (p=0.062, **Figure 3C**) or (p=0.85, **Figure 3D**), respectively. However, the comparison of 11A1-labeled pericapillary spaces revealed significantly less labeled spaces in the *APOE* ε3/ε4 genotype (p<0.0001, **Figure 3E**). Of note, in *APOE* ε*3/*ε*4* individuals the decrease in the percentage of 11A1-labeled pericapillary spaces became manifest in their 30s, whereas this change appeared after 45 years of age in *APOE* ε3/ε3 individuals (**Figure 3F**). Concomitant with the decrease in pericapillary labeling, we observed a significant increase in extracellular 11A1-labeled particles in *APOE* ε3/ε4 individuals compared to *APOE* ε3/ε3 individuals (p=0.0048, **Figure 3G**). Notably, extracellular 11A1 IR in *APOE* ε*3/*ε*4* individuals increased steadily from their early 30s, whereas in *APOE* ε*3/*ε*3* individuals, it begins to rise after age 45 years (**Figure 3H**).

**Figure 3.**
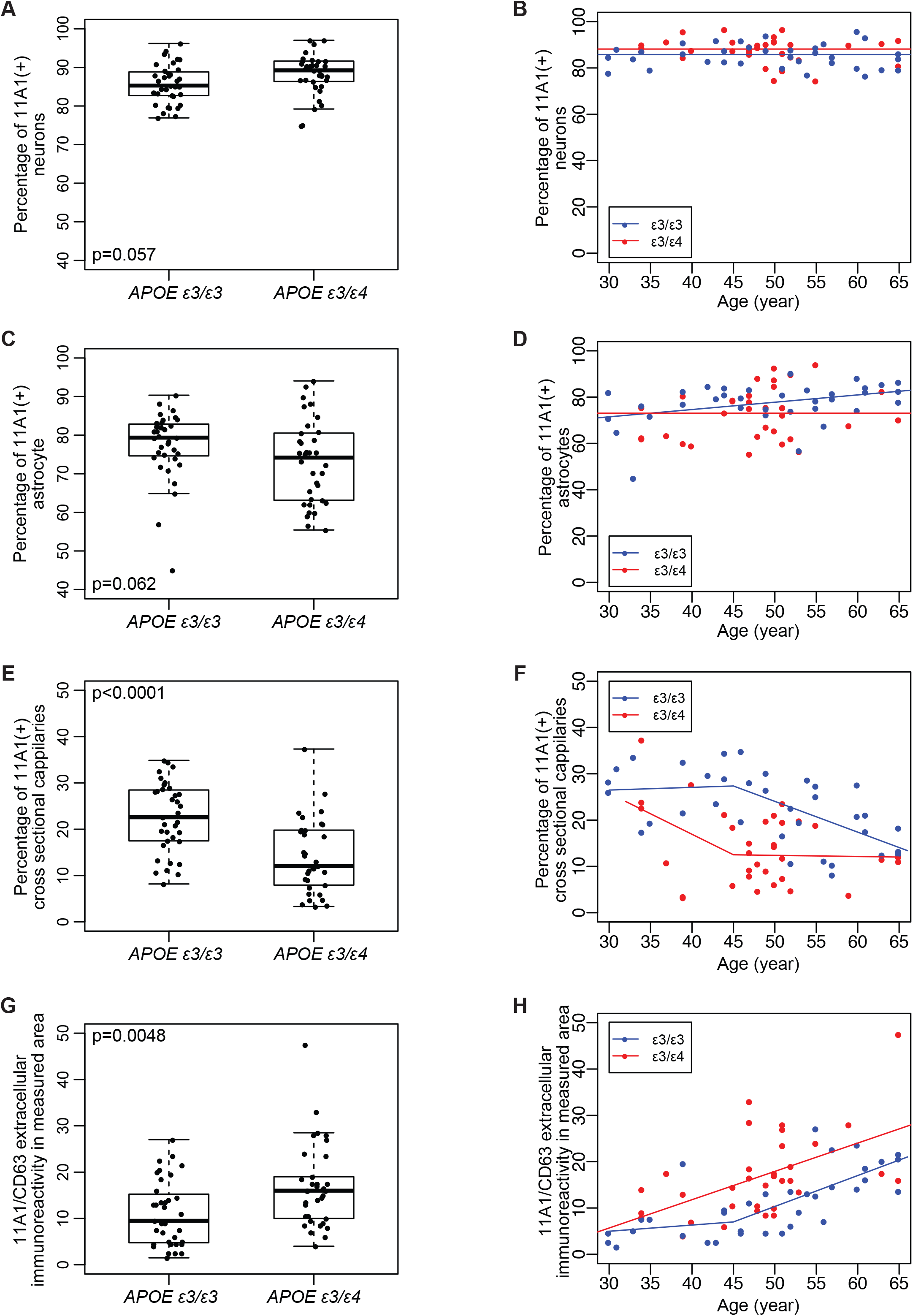
Comparison of 11A1 immunoreactivity between *APOE* ε*3/*ε*3* and *APOE* ε*3/*ε*4* individuals. Box and whisker plot of each value from *APOE* ε*3/*ε*4* individuals in **Figure 2** is compared with the results from *APOE* ε*3/*ε*3* individuals, **(A)** MAP2-, **(C)** ALDH1L1-, **(E)** Collagen IV-, and **(G)** CD63-labeled 11A1 immunoreactivity, respectively. No significant differences were observed in 11A1 IR in neurons (p=0.057) or astrocytes (p=0.062) (**A, C**). Pericapillary 11A1 IR was significantly decreased in *APOE* ε*3/*ε*4* individuals (p<0.0001) (**E**), and extracellular 11A1 IR was significantly increased in *APOE* ε*3/*ε*4* individuals (p=0.0048) (**G**). Student’s t-test was used to obtain p values. **(B, D, F, H)** Merged scatter plots of *APOE* ε*3/*ε*3* and *APOE* ε*3/*ε*4* individuals. Red is the data from *APOE* ε*3/*ε*4* individuals in **Figure 2**. Blue is from *APOE* ε*3/*ε*3* individuals. **(B)**MAP2-, **(D)** ALDH1L1-, **(F)** Collagen IV-, and **(H)** CD63-labeled 11A1 immunoreactivity, respectively.

### 3.5 Total area, size, and number of extracellular 11A1 immunoreactive Aβ aggregates (>3µm)

Based on our observations above, we hypothesized that the *APOE* ε*4* allele promotes Aβ aggregation preceding plaque formation^15^. To test our hypothesis, we examined extracellular Aβ aggregates, defined as structures displaying 11A1 and CD63 co-localization, measuring ≥3µm in diameter, and located at a distance ≥5µm from the edge of DAPI nuclear signal. We counted the Aβ aggregates in standard microscopic fields (0.407 mm^2^) and measured their individual and total areas in *APOE* ε*3/*ε*3* and *APOE* ε*3/*ε*4* individuals as described in eMethods. **Figure 4A** shows examples of aggregates of different sizes. The total area of the aggregates per image (area of image=0.407mm^2^) increased significantly with age in both of genotypes, *APOE* ε*3/*ε*3* (p<0.0001, r=0.66) (**Figure 4B**, blue) and *APOE* ε*3/*ε*4* (p<0.0001, r=0.69) (**Figure 4B**, red). The rates of increase were not different between genotypes (p=0.22, **Figure 4B**). However, the average total area value was significantly higher in *APOE* ε*3/*ε*4* individuals (p=0.0056) (**Figure 4C**). We further assessed the contribution of the size and number of individual Aβ aggregates to this age-related change. The average size of aggregates in both genotypes does not display any age-related changes (APOE ε3/ε3: p=0.45, r=-0.13, APOE ε3/ε4: p=0.74, r=-0.06, **Figure 4D**). After adjusting for age, the average size of aggregates in APOE ε3/ε3 is not significantly different from that in APOE ε3/ε4 (p=0.15, **Figure 4E**). As shown in **Figure 4F**, the number of Aβ aggregates increases significantly with age in both genotypes (*APOE* ε*3/*ε*3*: p<0.0001, r=0.67, *APOE* ε*3/*ε*4*: p=0.0005, r=0.57), but the rates of increase are not different between genotypes (p=0.41, **Figure 4F**). After adjusting for age, the average number of aggregates per area in APOE ε3/ε3 is significantly less compared with APOE ε3/ε4 (p = 0.0019, **Figure 4G**).

**Figure 4.**
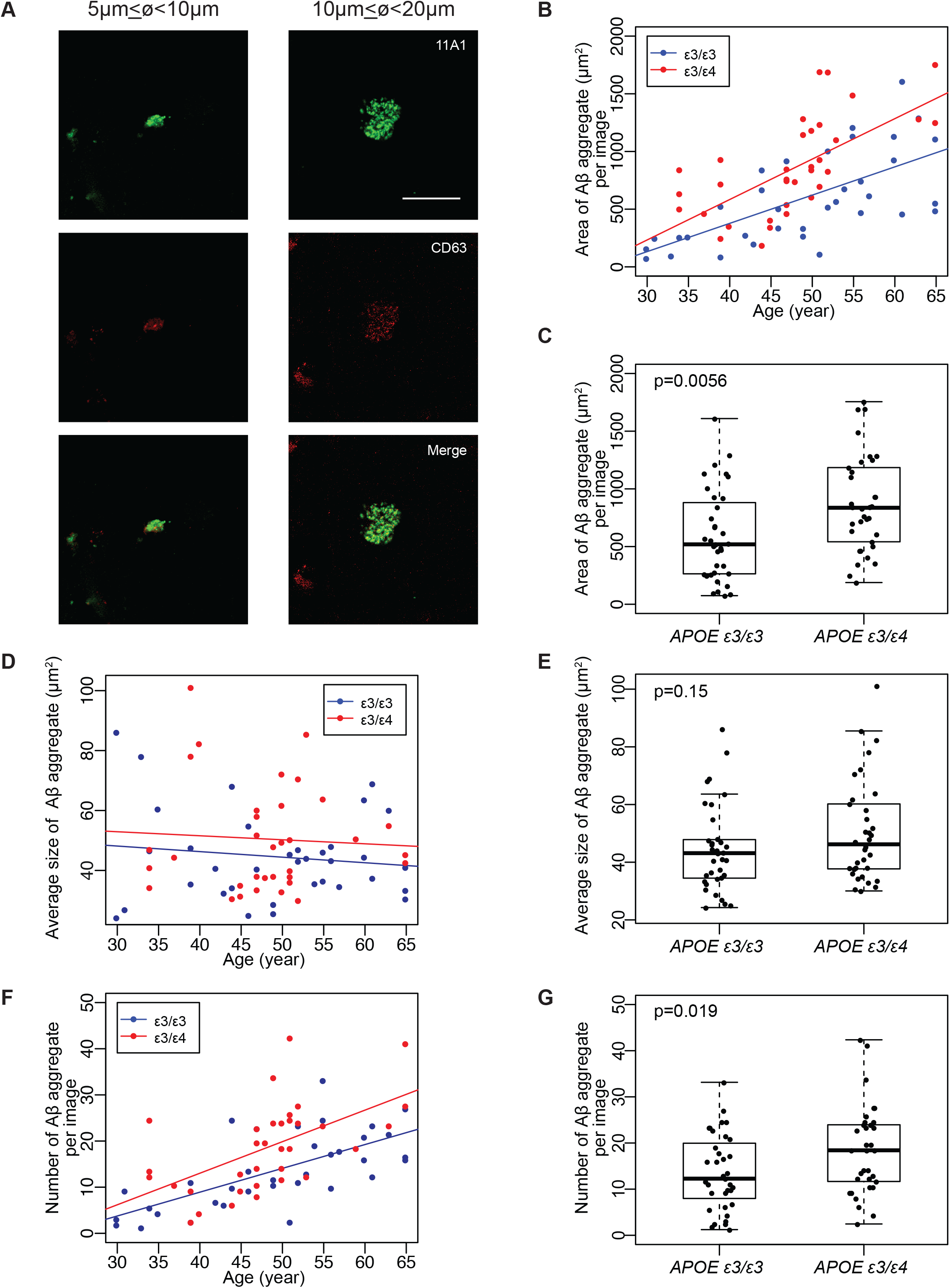
Amyloid Beta aggregates in the cortex. **(A)** Representative images of different sizes of Aβ aggregates. Approximate diameter of aggregates is indicated above the images. Extracellular Aβ aggregates were defined by colocalization of 11A1 and CD63 immunoreactivity. *Scale bar* 20µm applies to all images. **(B, D, F)** Scatter plot of average area (B), size (D) or number (F) of Aβ aggregates per image (0.407mm^2^) of each genotype, *APOE* ε*3/*ε*3* individuals (red) and *APOE* ε*3/*ε*4* individuals (blue). **(C, E, G)** Age-adjusted comparison of average area (C), size (E) or number (G) of Aβ aggregates between *APOE* ε*3/*ε*4* individuals and of *APOE* ε*3/*ε*3* individuals. **(B)** Linear regression analysis shows total area of Aβ aggregates increases by age in both *APOE* ε*3/*ε*3* (p<0.0001, r=0.66) and *APOE* ε*3/*ε*4* (p<0.0001, r=0.69). The interaction term is not statistically significant (p=0.22) in the regression model. **(C)** The average value in *APOE* ε*3/*ε*4* individuals is significantly higher than that in *APOE* ε*3/*ε*3* individuals (p=0.0056). **(D)** Linear regression analysis shows size of Aβ aggregates increases by age in both *APOE* ε*3/*ε*3* (p=0.45, r=-0.13) and *APOE* ε*3/*ε*4* (p=0.74, r=-0.06). **(E)** The average value of aggregate size in *APOE* ε*3/*ε*4* individuals is not significantly different from that in *APOE* ε*3/*ε*3* individuals (p=0.15). **(F)** Linear regression analysis shows the number of Aβ aggregates increases by age in both *APOE* ε*3/*ε*3* (p<0.0001, r=0.67) and *APOE* ε*3/*ε*4* (p<0.0001, r=0.57). The interaction term is not statistically significant (p=0.41) in the regression model. **(G)** The average value of Aβ aggregates number in *APOE* ε*3/*ε*4* individuals is significantly higher than that in *APOE* ε*3/*ε*3* individuals (p=0.0019).

## 4 DISCUSSION

The first aim of this study was to examine the dynamics of Aβ in the inferior parietal cortex of *APOE* ε*3/*ε*4* individuals 34 to 65 years of age, whose brains were free of Aβ plaques, and compare them to our previous observations in *APOE* ε*3/*ε*3* individuals^9^. We observed that the number of Aβ extracellular particles, co-labeled with 11A1/CD63, increased steadily with age at the same time that the presence of Aβ particles in pericapillary spaces declined. Notably, these changes appeared approximately a decade earlier in the *APOE* ε*3/*ε*4* individuals. The implication of these observations is that the glymphatic clearance of Aβ from the brain decreases with age and that the presence of *APOE* ε*4* allele further enhances this decline.

The specificity of the 11A1 Aβ antibody used in this study is crucial for the validity of our observations. Thus, we present new biophysical and immunological data on this reagent. Previously published results document that 11A1 recognizes Aβ in plaques and blood vessels of AD brains with the same pattern than 6E10 and 4G8 but does not react with APP^9^. Our new crystallographic study of the 11A1-Aβ immune complex further refines the immunoreactivity of 11A1 identifying its epitope as the Aβ amino acid sequence 10-14, located between the epitopes of 6E10 (aa6-10) and 4G8 (aa17-24)^16^. This new evidence strengthens the validity of 11A1 for immunohistochemical studies of Aβ in healthy and AD brains.

Aβ is cleared from the brain through several mechanisms and pathways including extracellular proteolytic degradation^17^, cellular degradation by neurons, astrocytes and microglia^18^, and by the cerebrovascular system. This latter system includes clearance through the blood-brain-barrier (BBB) and interstitial fluid (ISF) glymphatic drainage pathway^19,20^. Our observations of Aβ in pericapillary spaces are germane to the glymphatic system, a recently recognized network of perivascular pathways in the brain that allows the exchange of interstitial fluid (ISF) and cerebrospinal fluid (CSF) and the elimination of soluble proteins and metabolites, including Aβ, from the brain parenchyma. This system has three components: CSF influx pathway along the Virchow-Robin spaces that surround penetrating arterioles; the pericapillary space where the mixing of CSF and ISF takes place; and the efflux of CSF/IF along perivenular spaces that eventually drain into the cervical lymphatic nodes^6,19,21^. At the core of the glymphatic system is the pericapillary space lined by AQP4 expressing astrocytic endfeet, which are critical for the clearance of metabolites from the interstitial compartment and brain parenchyma. Our finding of decreased Aβ in pericapillary spaces concomitant with increased parenchymal Aβ strongly suggests an impairment of the passage of Aβ from the ISF into the pericapillary space.

Our observation that the clearance of Aβ from the human inferior parietal cortex declines with age is in keeping with observations of impaired glymphatic clearance in aged animals, which in old mice reaches 80% to 90% impairment compared to young animals^22^. This impairment has been attributed to proliferation of reactive astrocytes^23^, loss of vascular polarization of AQP4, and decline in CSF production^22^. Furthermore, a recent report described an age-associated decline in the drainage of CSF into the meningeal lymphatic vessels^21^. These abnormalities may render the aging brain vulnerable to Aβ accumulation in Alzheimer’s disease and amyloid angiopathy^11,24^.

Our findings are also consistent with observations that the human *APOE* isoforms differentially regulate the clearance of Aβ from the brain^25^. In that study, the mean concentration of CSF Aβ42 was significantly lower in *APOE* ε*4* carriers compared to *APOE* ε*3/*ε*3* individuals. This difference likely reflects a change in the equilibrium between the brain and CSF pools of Aβ. Even more relevant to our observations, that study showed that *APOE* isoform-dependent differences in soluble Aβ concentration and clearance from the brain are present before the onset of Aβ deposition. We observed a similar pattern of age-dependent decline in Aβ perivascular clearance and increase in parenchymal Aβ/CD63 particles modulated by *APOE* genotype preceding the deposition of Aβ in brain parenchyma or vessels. The interactions between Aβ and APOE are complex. These molecules interact with each other and share common receptors^15^. Both APOE and its receptors appear to regulate the clearance of Aβ through the blood-brain-barrier (BBB). For instance, LDL Receptor Related Protein 1 (LRP1) plays a key role in intracellular signaling and endocytosis. Whereas LRP1 enhances the Aβ clearance at the BBB, APOE works as a ligand of LRP1 and impairs the Aβ clearance^26^. Aβ and APOE form complexes that are cleared at the BBB by the LRP1 and VLDLR with the clearance rates being faster for Aβ-APOE3 or -APOE2 complexes than for Aβ-APOE4 complexes^27^. Although not known whether the APOE receptors that regulate the clearance of Aβ via the BBB are operative in the glymphatic system, it has been reported that the genetic ablation of *Aqp4* in APP/PS1 mice leads to enhanced Aβ accumulation in the brain of these animals^28^.

Our study has several caveats and limitations inherent to working with postmortem human tissues that include difficulty to evaluate variability in pre- and postmortem circumstances. The origin of the samples is from a forensic institution and as such is not representative of the population at large. The lack of cognitive information and of biochemical determination of Aβ in the CSF and brain tissues are also a relative weaknesses of our study.

In conclusion, our observations reveal a decline of Aβ clearance through a compartment of the glymphatic system starting about a decade earlier in *APOE* ε*3/*ε*4* than in *APOE* ε*3/*ε*3* individuals. This finding suggests that the timing for interventions to prevent Aβ accumulation in the brain and the development of AD should be contemplated as early as in the fourth and fifth decades of life and stratified by *APOE* genotype.

## ACKNOWLEDGEMENT

We would like to thank the staff members of Photon Factory for their help with X-ray data collections with synchrotron radiation. We also thank Dr. Masahiro Maeda (Immuno-Biological Laboratories Co, Ltd.) for providing Fab domain of 11A1 antibody for crystal structure analysis and Dr. Masahiro Fujihashi, Department of Chemistry, Graduate School of Science, Kyoto University, for support in the analysis of the X-ray crystal structure of 11A1 Fab-hapten peptide. We thank Ms. Karen Fisher for editorial assistance with the manuscript and our student members, Mr. Collin English, Mr. Tomonori Matsushita, Mr. Junya Fujino, and Mr. Ian Augsburger for their great help.

Authors’ contributions: AS, YK and JCT designed the project, analyzed and interpreted experiments. AS designed and performed the immunohistochemistry and confocal imaging. OP, GLR and JCT performed brain autopsies and neuropathological examinations. GLR performed *APOE* genotyping. YI and KI synthesized hapten peptides. YI, KI and AK prepared co-crystals. AK and KM analyzed the crystal data. YA contributed to statistical analyses. LL and PS supervised autopsies at the OCME. AS and JCT drafted the manuscript. All authors discussed results, edited and approved the manuscript. YK and KI reviewed the manuscript.

