## Supplementary Information for "*APOE ε4* allele advances the age-dependent decline of amyloid β clearance in the human cortex"

### A. 1 | SUPPLEMENTARY METHODS

#### A. 1.1 | Autopsy and Neuropathological examination

Following external examination and weighing, brains were cut in coronal slabs, diagnostic tissue blocks dissected, and the remainder tissues were frozen. For microscopic examination, tissues from the superior frontal gyrus, middle and superior temporal gyri, inferior parietal cortex, occipital cortex, amygdala, hippocampus and entorhinal cortex, midbrain, pons, medulla, and cerebellum were dissected in all brains. Tissues were fixed in 10% buffered formaldehyde, processed, embedded in paraffin, and cut at 10μm thickness. All sections were stained with H&E and screened for AD lesions using silver stains (Hirano method) [1], and immunostained for amyloid β (Aβ) (6E10) and phosphorylated tau (PHF-1). From a cohort of 523 autopsy brains from subjects 30-65 years of age, we selected brains free of Aβ plaques and tau pathology in the neocortex, and chose those with *APOE ε3/ε4* genotypes. In this way, we identified 34 brains suitable for the present study. All immunostainings were conducted in tissue sections from the inferior parietal cortex. We chose this region based on our previous neuropathological observations in this cohort [2] which demonstrated that the early deposition of Aβ is more common in the neocortex, including the inferior parietal cortex, than in the hippocampus. Also, the inferior parietal cortex is one of the cortical regions examined for the standardized neuropathologic assessment of AD according to CERAD guidelines[3]. The demography and causes of death of subjects examined in this study are listed in **Table A. 1**. Genotyping for *APOE* was performed using the method of Hixon and Vernier[4].

#### A. 1.2 | Crystallographic study of the 11A1 antibody

Fab of the 11A1 antibody was prepared from the papain digests and purified by gel filtration using Superdex 200 column (GE Healthcare) with PBS buffer. The purified Fab proteins were dialyzed against PBS buffer, after absorption of Fc fragments with Protein A Sepharose, and concentrated up to *ca.* 1 mg/mL, to which a hapten peptide (E22P-Aβ10-34) was added so as to adjust the molar ratio of the Fab and the peptide to 1:3. The mixture was further concentrated up to *ca.* 10 mg/mL using the Amicon centrifugation kit (Amicon Ultra 10 K, 14,000 *g*, 293 K, 7 min). Crystallization conditions were screened with the sitting-drop vapor diffusion method using commercial crystallization kits at 293 K. The crystals of 11A1-peptide complex were obtained from a 1:1 mixture of the protein solution and precipitant solution [25% (w/v) PEG3,350, and 0.2 M NaCl in 0.1 M Tris-Cl buffer (pH 8.5)]. The crystals belong to the space group $P2_1$ with cell dimensions of $a$=40.6 Å, $b$=123.8 Å, $c$=45.7 Å, and $\beta$=99.7˚.

Before the data collection, crystals were soaked for 15 - 20 sec. in precipitant solutions containing 20% ethylene glycol as cryogenic solutions, and then frozen in liquid $N_2$. X-ray diffraction work was carried out under a nitrogen-gas stream at 100 K using synchrotron radiation with a Pilatus 2M detector on the ARNE3A beamline at Photon Factory (Tsukuba, Japan). The diffraction data were processed and scaled with the XDS package[5] and Aimless program[6] from the CCP4 suite[7]. The structure of 11A1-Fab complex was solved at 1.75 Å resolution by the molecular replacement method using the program Molrep[8] from the CCP4[7] suite, with the coordinates of bactericidal antibody (PDBcode:1MNU)[9] as a search model. The models were manually constructed using the program Coot[10], and structure refinement was performed with the program REFMAC5[7,11]. The final model of 11A1-peptide consists of L-chain, H-chain, peptide (residue No. 10-14), 5 ethylene glycols, and 431 water molecules. The quality of the model was checked using the MolProbity server[12] at http://molprobity.biochem.duke.edu/. The data collection and the refinement statistics are summarized in **Table A. 5**.

Pictures of crystal structure were prepared using the program PyMOL (The PyMOL Molecular Graphics System, Version 1.2r3pre, Schrödinger, LLC) unless otherwise stated.

#### A. 1.3 | PDB references

The atomic coordinates and structure factors of 11A1-peptide complex have been deposited in the Worldwide Protein Data Bank, with the accession code of 7BXV.

**A. 1.4 | Immunohistochemistry with fluorescent probe**

All tissue sections were deparaffinized in xylene and rehydrated in 100% and 95% EtOH**.** To ascertain good preservation of tissues to be examined, we screened sections of the cerebellum with β-tubulin immunohistochemistry as previously described[13]. For cerebellum, antigen retrieval was performed in 1x Citrate buffer (pH6.0) (Abcam: ab93678) by boiling for 4 min. The samples from the inferior parietal cortex to be co-stained with the 11A1 antibody were incubated in 70% formic acid (Sigma: F0507) for 12 min at room temperature (RT) prior to antigen retrieval. After washing with tap water, antigen retrieval was performed in 1mM EDTA (pH8.0) (Invitrogen: 15575-038) by boiling for 4 min. Then, all samples were blocked in PBS (Phosphate Buffered Saline) with 5% normal goat serum and 0.2% Triton x-100 for 1 hr at RT. Primary antibodies were incubated in blocking buffer for 16h at 4°C. On the following day, samples were washed in PBS for 5 min x 3, then we applied Alexa fluor secondary antibodies in PBS with 0.5% Tween 20 and incubated for 1 hr at RT. Samples were washed in PBS once for 5 min, then 5μg/mL Hoechst 33258 in PBS were applied and incubated for 20 min at RT. Samples were washed in PBS once for 5 min. For quenching lipofuscin autofluorescence, we used TrueBlackTM Lipofuscin Autofluorescence Quencher (Biotium: 23007) diluted 1/40 in 70% EtOH, applied to the samples and incubated for 50 seconds at RT. To facilitate the TrueBlack reaction, samples were constantly swirled by hand during the incubation. Then, samples were washed in PBS for 5 min x 3 and coverslipped using ProLongTM Gold Antifade reagent (Invitrogen: P36930). Stained tissue sections were kept at 4°C at least 2 days before imaging. Immunofluorescent images were taken on a Zeiss LSM 700 confocal microscope in the Microscope Facility of the Johns Hopkins School of Medicine.

**A. 1.5 | Imaging and data analysis**

**A. 1.5.1 | Quantitation of 11A1 labeled neurons, astrocytes and pericapillary spaces**

All images were captured from layer V of the inferior parietal cortex. The cortical layer V was identified as the lamina of pyramidal neurons closest to the gray-white matter junction. To calculate the percentage of 11A1 labeled cortical neurons and protoplasmic astrocytes, first we immunostained tissue with 11A1, MAP2 and ALDH1L1 antibodies and captured two montage images, one at the crown of the gyrus, and the other midway between the crown of the gyrus and the bottom of the sulcus. Each montage consisted of a 3 x 3 panel of images captured with a 20x objective lens encompassing an area of 0.910mm$^2$. The number of neurons and astrocytes with and without 11A1 signals in the measured field were counted. The number of 11A1 labeled neurons and astrocytes was divided by total number neurons and astrocytes to calculate the percentage of cells with 11A1 signal. We stained an adjacent tissue section with 11A1 and Collagen IV antibodies to calculate the percentage of 11A1 labeled pericapillary spaces using the same imaging approach employed for neurons and astrocytes. For the quantification of pericapillary spaces labeled with 11A1 signals, we selected only cross-sectional pericapillary spaces with diameters ≤20μm to avoid interpretation problems arising from measuring longitudinal profiles. The number of pericapillary spaces with and without 11A1 signal in the measured field were counted and used to calculate the percentage of 11A1 labeled pericapillary spaces. The calculated value from the two montage images in each sample was averaged for the final analysis.

**A. 1.5.2 | Quantitation of Aβ parenchymal extracellular particles and aggregates**

We also observed parenchymal extracellular particles and aggregates that co-labelled for 11A1 and CD63. We counted the number of these particles and measured their sizes. To this end, we set as criterion for identification of extracellular 11A1 immunoreactive particles and aggregates at a distance ≥ 5µm from DAPI signal. As for size, particles are ≤ 1µm and aggregates are ≥3µm in diameter.

First, we immunostained the tissue with the 11A1 and CD63 antibodies followed with DAPI incubation. Then, we captured two montage images of the cortical layer V as described above. These montages consisted of a 4 x 4 panel of images captured with a 40x objective lens encompassing an area of 0.407mm$^2$. The number of 11A1/CD63 particles was measured within these areas. The calculated value from two montage images in each sample was averaged for the final analysis. We used Image J for the measurement of the size of 11A1/CD63 immunoreactive aggregate. For the final analysis, we calculated the average value from the two montage images for each sample. Representative images were obtained under 63x objective lens (**Figure 2**, **Figure 4** and **Figure A. 1**).

Figure A.1

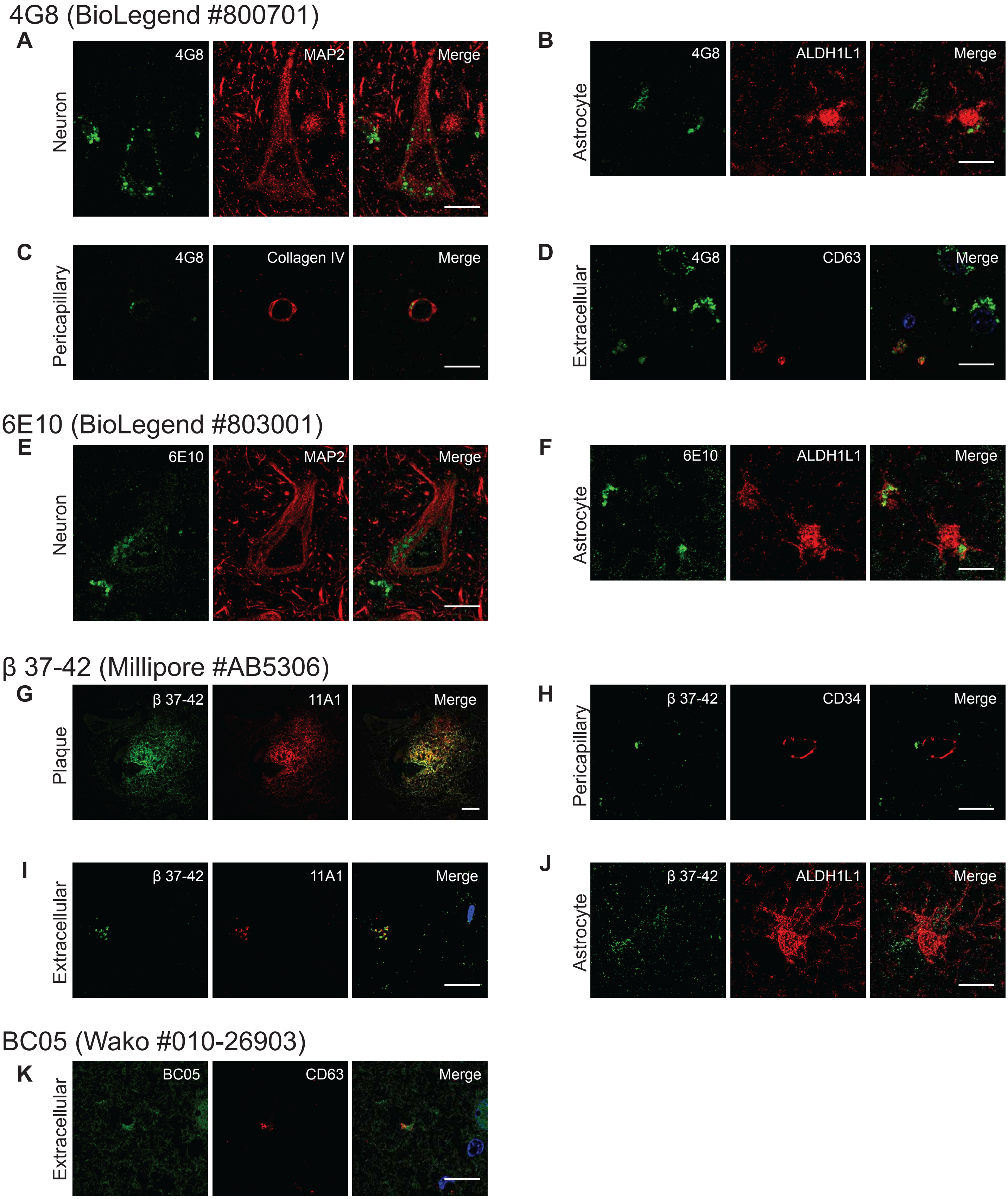
4G8 (BioLegend #800701)
A
Neuron
4G8
MAP2
Merge
B
Astrocyte
4G8
ALDH1L1
Merge
C
Pericapillary
4G8
Collagen IV
Merge
D
Extracellular
4G8
CD63
Merge
6E10 (BioLegend #803001)
E
Neuron
6E10
MAP2
Merge
F
Astrocyte
6E10
ALDH1L1
Merge
β 37-42 (Millipore #AB5306)
G
Plaque
β 37-42
11A1
Merge
H
Pericapillary
β 37-42
CD34
Merge
I
Extracellular
β 37-42
11A1
Merge
J
Astrocyte
β 37-42
ALDH1L1
Merge
BC05 (Wako #010-26903)
K
Extracellular
BC05
CD63
Merge

**Figure A. 1. Amyloid β immunoreactivity by commercially available miscellaneous antibody.** Representative images of Amyloid β (Aβ) antibodies co-stained with another antibodies including molecular markers; **(A)** 4G8 and MAP2, **(B)** 4G8 and ALDH1L1, **(C)** 4G8 and Collagen IV, **(D)** 4G8 and CD63, **(E)** 6E10 and MAP2, **(F)** 6E10 and ALDH1L1, **(G)** β 37-42 and 11A1 for Aβ plaque, **(H)** β 37-42 and CD34, **(I)** β 37-42 and 11A1 for extracellular vesicle, **(J)** β 37-42 and ALDH1L1, and **(D)** BC05 and CD63, respectively. White arrows indicate the positions of immunoreactivity. *Scale bars*, 5μm.

Table A. 1: Demography and cause of Death of subjects

| Sample # | Sex | Age | Race | Cause of death |
|---|---|---|---|---|
| 1 | M | 34 | Caucasian | Combined drug intoxication |
| 2 | M | 34 | African American | Combined drug intoxication |
| 3 | F | 34 | Caucasian | Pulmonary Thromboembolism |
| 4 | M | 37 | Caucasian | Combined drug intoxication |
| 5 | F | 39 | Caucasian | Drug intoxication |
| 6 | M | 39 | Other | Multiple Injuries |
| 7 | F | 40 | Caucasian | Asphyxia |
| 8 | F | 44 | Caucasian | Combined drug intoxication |
| 9 | F | 45 | Caucasian | Cardiovascular Disease |
| 10 | F | 45 | Caucasian | Combined drug intoxication |
| 11 | F | 47 | Caucasian | Drug intoxication |
| 12 | M | 47 | Caucasian | Megacolon |
| 13 | F | 47 | Caucasian | Combined drug intoxication |
| 14 | M | 47 | African American | Combined drug intoxication |
| 15 | M | 47 | Caucasian | Cardiovascular Disease |
| 16 | M | 48 | Caucasian | Complications of Chronic Alcohol Use |
| 17 | F | 49 | Caucasian | Combined drug intoxication |
| 18 | M | 49 | Caucasian | Acute Coronary Artery Thrombosis |
| 19 | M | 50 | Caucasian | Diabetic Ketoacidosis |
| 20 | F | 50 | African American | Hyperosmolar Hyperglycemic Non-Ketotic Syndrome |
| 21 | M | 50 | Caucasian | Combined drug intoxication |
| 22 | M | 50 | Caucasian | Cardiovascular Disease |
| 23 | M | 51 | Caucasian | Cardiovascular Disease |
| 24 | M | 51 | Caucasian | Cardiovascular Disease |
| 25 | M | 51 | Caucasian | Asphyxia |
| 26 | F | 51 | Caucasian | Cardiovascular Disease |
| 27 | M | 52 | African American | Combined drug intoxication |
| 28 | M | 52 | Caucasian | Cardiovascular Disease |
| 29 | M | 53 | African American | Complications of embedded esophageal foreign body |
| 30 | F | 55 | African American | Asphyxia |
| 31 | M | 59 | Caucasian | Cardiovascular Disease |
| 32 | M | 63 | Caucasian | Cardiovascular Disease |
| 33 | M | 65 | Caucasian | Multiple Injuries |
| 34 | M | 65 | African American | Multiple Injuries |

Table A. 2: List of primary antibodies used in this study

| Antibody | Company | Catalog # | Host | Dilution | Target |
|---|---|---|---|---|---|
| 11A1 | IBL | 10379 | Mouse | 1/50 | Amyloid β |
| 4G8 | BioLegend | 800701 | Mouse | 1/250 | Amyloid β |
| 6E10 | BioLegend | 803001 | Mouse | 1/250 | Amyloid β |
| β 37-42 | Millipore | AB5306 | Rabbit | 1/100 | Amyloid β |
| BC05 | Wako | 010-26903 | Mouse | 1/250 | Amyloid β |
| ALDH1L1 | abcam | ab177463 | Rabbit | 1/100 | Astrocyte |
| CD34 | Beckman Coulter | IMO0786 | Mouse | 1/500 | small blood vessels/capillaries |
| CD63 | abcam | ab134045 | Rabbit | 1/200 | Tetraspanin family protein |
| Class III β-tubulin | abcam | ab18207 | Rabbit | 1/1000 | Neuronal β-tubulin |
| Collagen IV | abcam | ab6586 | Rabbit | 1/500 | Collagen in perivascular space |
| MAP2 | abcam | ab92434 | Chicken | 1/1000 | Neuron |

Table A. 3: List of secondary antibodies used in this study

| Antibody | Company | Catalog # | Host | Dilution |
|---|---|---|---|---|
| Anti-Chicken IgY H&L (Alexa Fluor 568) | Abcam | ab175711 | Goat | 1/400 |
| Anti-Mouse IgG H&L (Alexa Fluor 488) | Abcam | ab150117 | Goat | 1/400 |
| Anti-Mouse IgG H&L (Alexa Fluor 568) | Abcam | ab175701 | Goat | 1/400 |
| Anti-Rabbit IgG H&L (Alexa Fluor 488) | Abcam | ab150081 | Goat | 1/400 |
| Anti-Rabbit IgG H&L (Alexa Fluor 568) | Abcam | ab175696 | Goat | 1/400 |
| Anti-Rabbit IgG H&L (Alexa Fluor 647) | Abcam | ab150083 | Goat | 1/400 |

Table A. 4: Akaike Information Criterion (AIC) values

| | | AIC |
|---|---|---|
| MAP2 | Intercept 1 | 213.6 |
| | Linear age 2 | 214.3 |
| | Piecewise 3 | 216.3 |
| ALDH1L1 | Intercept 1 | 260.9 |
| | Linear age 2 | 261.2 |
| | Piecewise 3 | 261.6 |
| Collagen IV | Intercept 1 | 233.8 |
| | Linear age 2 | 232.2 |
| | Piecewise 3 | 231.8 |
| CD63 | Intercept 1 | 248.9 |
| | Linear age 2 | 239.5 |
| | Piecewise 3 | 241 |

Red indicates the best fitted model.

Table A.5: Data collection and refinement

| | |
|---|---|
| Data collection statistics | |
| Protein | 11A1-peptide |
| PDB code | 7BXV |
| Wavelength | 1.000 |
| Space group | $P2_1$ |
| Cell dimensions (Å,˚) | |
| *a* | 40.6 |
| *b* | 123.8 |
| *c* | 45.7 |
| $\beta$ | 99.7 |
| No. of molecules per asymmetric unit | 1 |
| Resolution (Å) | 1.75 |
| Measurements | 304,239 |
| Unique reflections | 44,761 |
| $R_{merge}$[a] | 0.063 (0.135)[b] |
| Multiplicity | 6.8 (6.8) |
| $I/\sigma(I)$ | 21.5 (11.1) |
| Completeness (%) | 100.0 (99.8) |
| Overall B factor from Wilson plot (Å$^2$) | 11.6 |
| Refinement statistics | |
| Resolution range (Å) | 1.75 |
| No. of reflections | 42,474 |
| $R_{work}/R_{free}$[c,d] (%) | 0.166/0.194 |
| No. of atoms | |
| Fab/peptide/others | 3,317/48/451 |
| Average *B* factors | |
| Fab/peptide/others (Å$^2$) | 16.6/11.3/27.5 |
| R. m. s. d. from ideal geometry | |
| Bond length (Å) | 0.003 |
| Bond angles (˚) | 1.275 |
| Ramachandran plot | |
| Favored /outliers (%) | 98.4/0.0 |

[a] $R_{merge} = \Sigma (|I-<I>|)/\Sigma (I)$

[b] Numbers in parentheses refer to the highest-resolution shell, 1.78-1.75 Å.

[c] $R = \Sigma||F_{obs}|-|F_{calc}||/\Sigma|F_{obs}|$

[d] $R_{work}$ is calculated from a set of reflections in which 5% of the total reflections have been randomly omitted from the refinement and used to calculate $R_{free}$.
